# VICTOR: Genome-based Phylogeny and Classification of Prokaryotic Viruses

**DOI:** 10.1101/107862

**Authors:** Jan P. Meier-Kolthof, Markus Göker

## Abstract

Bacterial and archaeal viruses (“phages”) play an enormous role in global life cycles and have recently regained importance as therapeutic agents to fight serious infections by multi-resistant bacterial strains. Nevertheless, taxonomic classification of phages is up to now only insufficiently informed by genome sequencing. Despite thousands of publicly available phage genomes, it still needs to be investigated how this wealth of information can be used for the fast, universal and accurate classification of phages. The Genome BLAST Distance Phylogeny (GBDP) approach is a truly whole-genome method currently used for *in silico* DNA: DNA hybridization and phylogenetic inference from prokaryotic genomes. Based on the principles of phylogenetic systematics, we here established GBDP for phage phylogeny and classification, using the common subset of genome-sequenced and officially classified phages. Trees inferred with the best GBDP variants showed only few deviations from the official phage classification, which were uniformly due to incorrectly annotated GenBank entries. Except for low resolution at the family level, the majority of taxa was well supported as monophyletic. Clustering genome sequences with distance thresholds optimized for the agreement with the classification turned out to be phylogenetically reasonable. Accordingly modifying genera and species is taxonomically optional but would yield more uniform sequence divergence as well as stronger branch support. Analysing an expanded data set containing > 4000 phage genomes from public databases allowed for extrapolating regarding the number, composition and host specificity of future phage taxa. The selected methods are implemented in an easy-to-use web service “VICTOR” freely available at http://ggdc.dsmz.de/victor.php.

## Introduction

Viruses are ancient (Forterre 2006), infect organisms of all three domains of life and outnumber the cells of their hosts by one to two orders of magnitude (Suttle 2007). Viral impact on the global biogeochemical cycles and thus on life on earth is immense (Suttle 2007). Since prokaryotes are the most abundant organisms on this planet (Whitman et al. 1998), they are the most common hosts of viruses. Consequently, "phages" (bacterial and archaeal viruses) are most abundant biological entities, not only in the virosphere itself but on earth in general (Koonin et al. 2015). Even though the genetic diversity of phages is unmatched (Hendrix 2003; Kristensen et al. 2013), probably only a minor fraction of them are known (Rohwer 2003). In recent years, the interest in bacteriophages as therapeutic agents to fight serious bacterial infections caused by multi-resistant strains (Abedon et al. 2011) experienced a renaissance (Thacker 2003). Likewise, the role of viruses in aquatic ecosystems (Wommack and Colwell 2000), especially by infecting and lysing aquatic microorganisms, and their subsequent impact on life-cycle and evolution of groups such as *Rhodobacteraceae* (Simon et al. 2017), is still not fully understood.

A reliable classification of the ever increasing number of known phages (Ackermann 2007) is thus of utmost importance (Ackermann 2011). The International Committee on Taxonomy of Viruses (ICTV) (Wildy 1971) established rules for the naming and classification of viruses, represented by the International Code of Virus Classification and Nomenclature and published in its latest version in 2016 (Adams et al. 2013; Krupovic et al. 2016). Whereas viruses are taxonomically arranged by a variety of characteristics (Ackermann 2009), the affiliation of phages to the same family is currently possible despite a complete lack of DNA sequence relatedness (Nelson 2004). A stronger role of genome sequences in phage classification was recommended though (Krupovic et al. 2016; Simmonds et al. 2017). As of March 2015, the ICTV master species list recognizes phages assigned to 548 species, 104 genera and 18 families but a much larger number of phage genomes is available in public databases.

DNA: DNA hybridization experiments with phages were already conducted by Grimont and Grimont (1981), who found stable hybridization groups and proposed to define a phage species as a group of phages with “significant” genomic relatedness. This technique is not routinely applied anymore due to an emphasis on genomics (Ackermann 2009), and, in contrast to the classification of bacteria and archaea, bacteriophage taxonomy has not yet established firm distance or similarity thresholds for assigning phage genomes to a taxon of a given taxonomic rank (Krupovic et al. 2016). In microbiology, the 70% DNA: DNA hybridization threshold has been applied since decades as the ultimate criterion to separate species (De Ley 1970; Brenner 1973; Johnson 1973; Wayne et al. 1987). It was recently successfully replaced by bioinformatic techniques applied to partial or complete whole-genome sequences (Auch et al. 2010a; Meier-Kolthoff et al. 2013). Boundaries for the subspecies rank have also been suggested (Meier-Kolthoff et al. 2014b), based on the same kind of intergenomic distances calculated with the Genome BLAST Distance Phylogeny (GBDP) tool (Henz et al. 2005; Auch et al. 2010a; Auch et al. 2010b). These techniques have previously been made available as an easy-to-use web service (Auch et al. 2010a; Auch et al. 2010b).

Several sequence-based approaches were specifically designed to tackle phage classification (Rohwer and Edwards 2002) but sometimes addressing the genus and family level only (Lima-Mendez et al. 2008; Roux et al. 2015) or restricted to a single taxonomic rank within a particular virus family (Lavigne et al. 2008) or subfamily (Asare et al. 2015). Bacteriophages can be classified according to their inferred neck organization (Lopes et al. 2014) but this is applicable only to tailed phages. Some tools (Frazer et al. 2004; Ågren et al. 2012; Asare et al. 2015) were recently suggested by Krupovic et al. (2016) for phage taxonomy but were not primarily devised for this purpose and are thus not necessarily optimal. Clustering techniques were frequently applied but not necessarily optimized for phage classification, with the notable exceptions of Lima-Mendez et al. (2008) and Roux et al. (2015). Only few methods attempted to incorporate phylogenetic analyses but these either were based on a predefined selection of few genes (Asare et al. 2015) or inferred trees but no branch support (Rohwer and Edwards 2002). Methods using standard sequences alignment techniques (Lavigne et al. 2008; Ceyssens et al. 2011) only work if the viral genomes are collinear. Most of the suggested approaches were restricted to amino-acid sequences (Lavigne et al. 2008; Lima-Mendez et al. 2008; Ågren et al. 2012; Lopes et al. 2014; Roux et al. 2015), and methods based on gene content were seldom compared to those based on sequence similarity (Rohwer and Edwards 2002).

Thus a comprehensive optimization of methods for phage classification, covering both clustering techniques and phylogenetic inference (including branch support), nucleotide and amino-acid sequences, and gene-content based, sequence-similarity based and combined strategies, appears to be missing. GBDP is promising in this respect because of its variety of options for determining homologous regions between genomes with which it can be combined (Auch et al. 2010a; Meier-Kolthoff et al. 2013; Meier-Kolthoff et al. 2014a), its algorithms for correcting for paralogy (Henz et al. 2005), the variety of GBDP distance formulas that explore distinct genomic features (Auch et al. 2010a; Meier-Kolthoff et al. 2013; Meier-Kolthoff et al. 2014a), and the ability to infer phylogenetic trees including branch-support values based on resampling (Meier-Kolthoff et al. 2014a).

Indeed, clustering based on sequence or distance thresholds on the one hand and phylogenetic inference on the other hand may be in conflict with each other. When distance matrices deviate from ultrametricity (Swofford et al. 1996), which usually happens if the evolution of the underlying sequence data deviated from a molecular clock (Felsenstein 2004), clusters based on a given distance or similarity threshold might not correspond to monophyletic groups (Wood 1994). The principles of phylogenetic to systematics imply that the goal of taxonomic classification is to summarize the phylogeny of the organisms under study (Hennig 1965; Wiley and Lieberman 2011). Non-monophyletic taxa (Farris 1974) are in conflict with that goal (Hennig 1965; Wiley and Lieberman 2011); the ideal classification comprises only taxa that are statistically well supported as monophyletic (Vences et al. 2013). Whereas ultimately rather a problem of the data and not of the methods, the severity of taxonomic problems due to non-ultrametricity depends on the organisms under study, the characters collected from these organisms and the methods used to analyse those characters. For instance, when using intergenomic distances inferred with GBDP to assign archaea and bacteria to species, few issues caused by non-ultrametric data were found (Meier-Kolthoff et al. 2014c). The same holds for subspecies, provided the distance or similarity threshold (and clustering approach) is chosen for maximum cluster consistency (Meier-Kolthoff et al. 2014b).

To create an easy-to-use web service for phage phylogeny, in this study we optimize the combination of GBDP parameters, analyzed at both nucleotide and amino acid level, to yield as many phage taxa highly supported as monophyletic as possible, quantified as “taxon support”. The ICTV classification combined with accordingly taxonomically annotated phage genome sequences from public repositories is used as reference data set. As secondary criterion, we optimize GBDP settings, clustering algorithms and distance thresholds to yield the highest agreement with the ICTV assignment of the phage genomes to taxa at distinct ranks, quantified via the Modified Rand Index (MRI) as implemented in OPTSIL (Göker et al. 2009). The results are discussed regarding the proportion of taxa that could be kept as-is and the number of revisions needed when broadly implementing the standardized approach and other potentials and limitations of the suggested method, such as those ranks too high to be resolved any more using GBDP. As an example for applying the devised approach to phylogeny and classification, we analyse an expanded data set of virtually all publicly available phage genomes. The outcome is quantified regarding the number of overall, known and novel taxa and regarding the detectable host specificity. Our phylogenomic approach contributes to understanding the composition of the viral biosphere by means of its accurate phylogeny and classification.

## Results

### Optimal GBDP Settings

The reference data set included 610 phage genomes (Supplementary File S1). The total number of pairwise intergenomic distances were c. 4.5B (185,745 pairwise comparisons × 100 replicates × 240 parameter combinations). The average branch support ranged from 25 % to 97 % (median: 50 %) for the 120 amino-acid trees and from 2 % to 92 % (median: 42 %) for the 120 nucleotide trees. For applying OPTSIL to the genus rank, the set had to be reduced to 588 genomes (i.e. 22 phages without genus affiliation).

The biplot of the two most important principal components in Fig. 1 shows the relative directions and loadings of the six optimality criteria (Supplementary File S1). Apparently the majority of the criteria, including all taxon support objectives, indicate approximately the same optimal GBDP settings, whereas MRI values for species and family, respectively, point into distinct directions. Table 1 shows the GBDP settings that yielded the highest taxon support and among those the GBDP settings and according distance threshold (*T*) values that yielded the highest MRI values. Nucleotide and amino-acid sequences both favored the greedy-with-trimming algorithm (Meier-Kolthoff et al. 2014a) but required distinct distance formulas, BLAST+ (Camacho et al. 2009) word lengths and e-values; amino-acid sequences yielded (i) higher average branch support, (ii) marginally lower taxon support values for species and genera, (iii) a pronounced lower taxon support value for families, (iv) similarly high MRI values for species and genera and (v) a higher (but overall still comparatively low) MRI value for the family rank.

**Figure 1.**
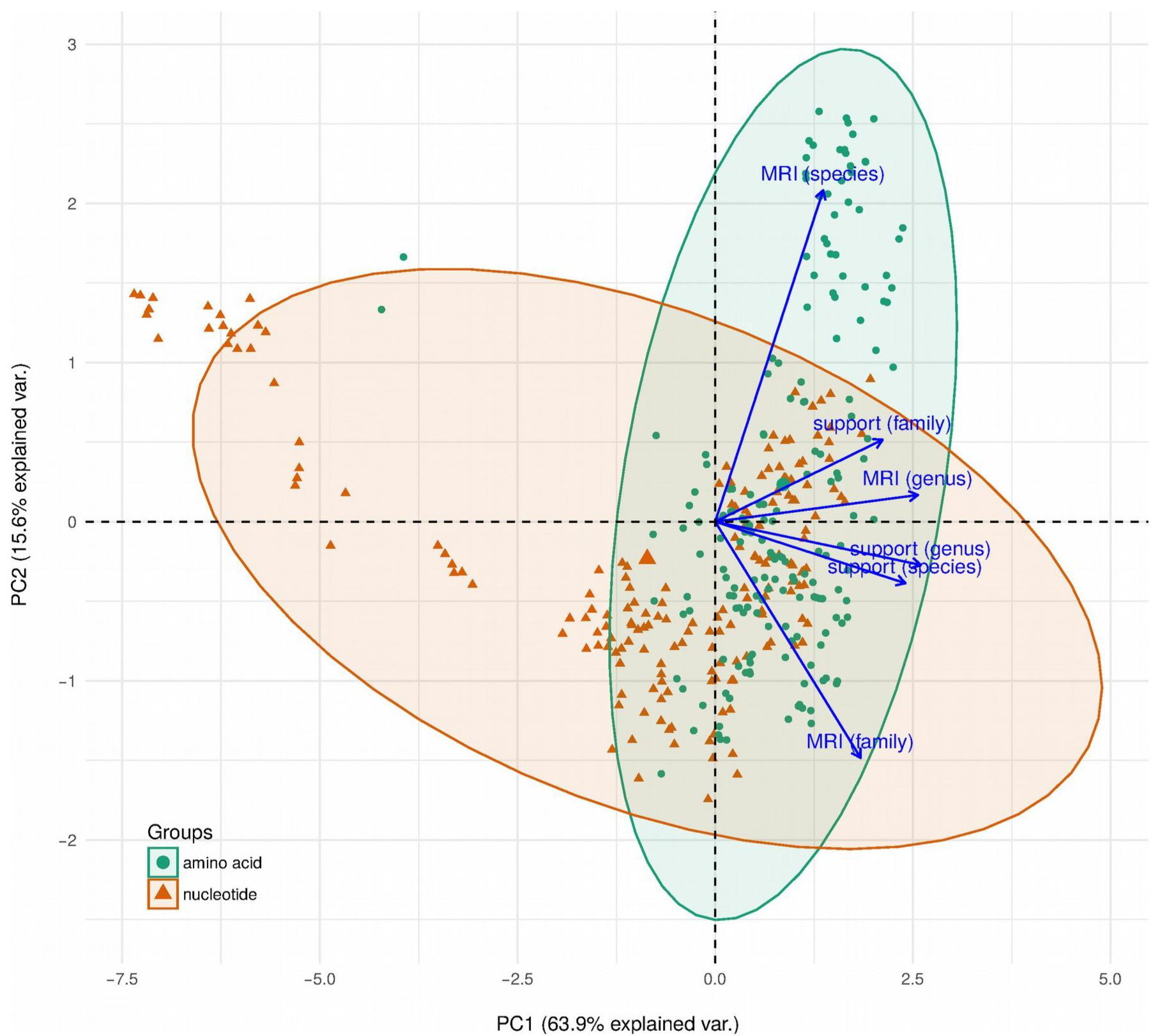
Relative performance of GBDP settings and optimality criteria. The two principal components (PC1 and PC2) that explained most of the variation (given in percent) in the data are displayed. Dots represent the individual GBDP settings, whereas arrows represent the loadings of the relevant variables and thus the performance of each combination of GBDP parameters. Normal probability ellipsoids are given separately for amino acid and nucleotide sequence and 95% confidence limits.

### Remaining non-monophyletic taxa

Fig. 2 shows the phylogenomic tree inferred from the amino-acid sequences under the optimal GBDP settings (Table 1). It is also contained in Supplementary File S3 in linear shape to allow for more detailed analysis; the according nucleotide tree is contained in Supplementary File S4. Families did not usually appear as monophyletic, but did not induce significant conflict either, due to the low resolution of the backbone of the tree, especially in the case of the nucleotide sequences. In contrast, particularly the genera were highly supported as monophyletic with only few exceptions (Supplementary File S1). The distribution of the positive and negative support values for each taxon (Fig. 3) indeed indicated only a handful of well-supported conflicts. These were uniformly due to problematic assignments of names rather than due to the GBDP tree.

**Figure 2.**
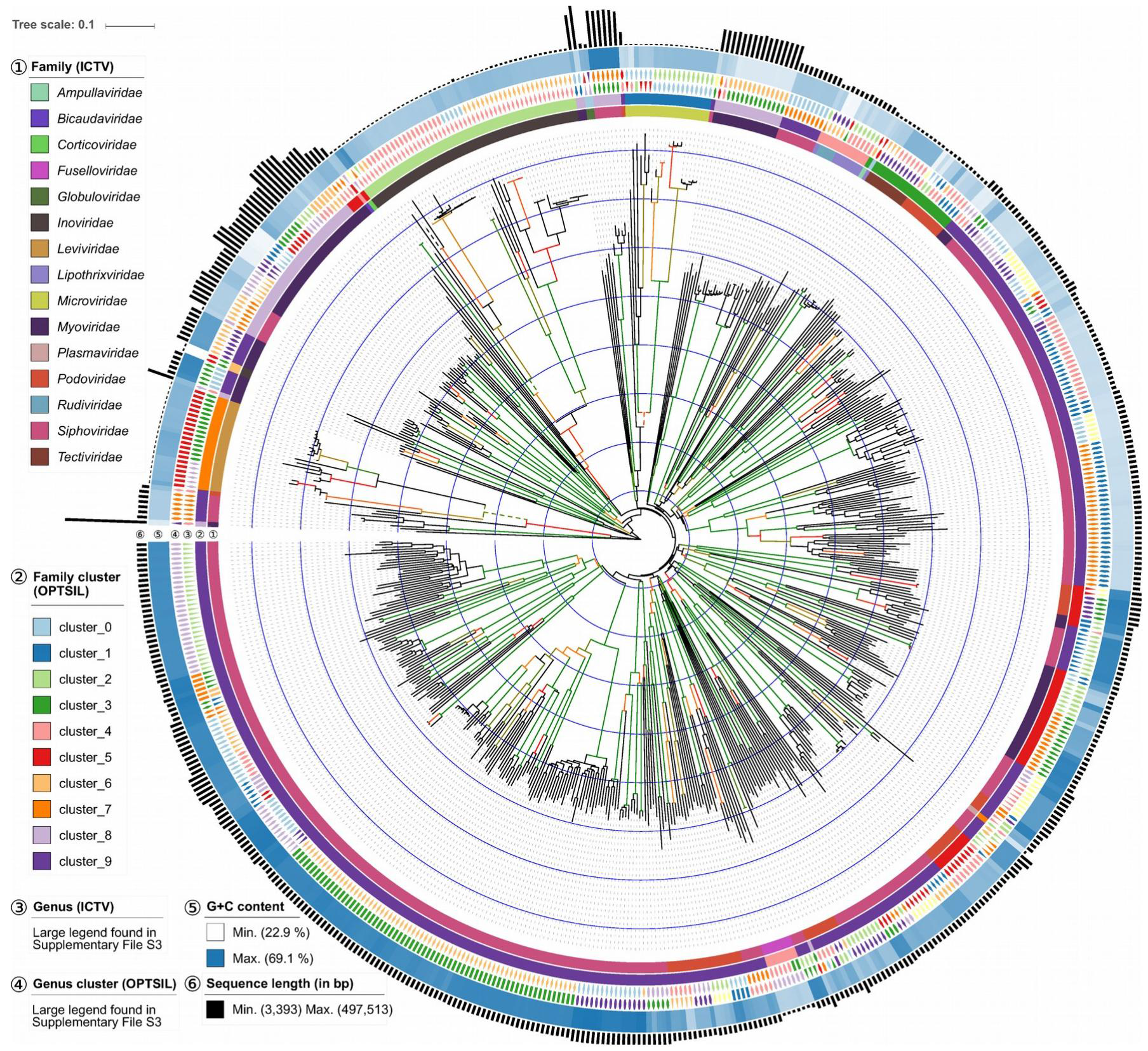
Phage phylogeny inferred from the amino-acid sequences contained in the reference date set. Four dashed branches were downscaled by a factor of four to ease visualization of the tree. Genome size, genomic G+C content, ICTV genera and families as well as clusters derived from the ICTV genera and families are shown next to the tips within circles 1-6. For both the tree and the clusterings the optimal GBDP settings (Table 1) were used. Branch support ≥ 60 % is shown as a colour gradient from red to green; terminal branches and branches with support < 60 % are black. A linear visualization is provided in Supplement S3.

For instance, genus *L5likevirus* was supported as non-monophyletic with 100 % support because of the positioning of *Mycobacterium phage Ta17a*. Genome sizes and G+C content values confirm that *Ta17a*, which was not included in the original description of *L5likevirus* (Hatfull and Sarkis 1993), deviates from the majority of the representatives of the genus and thus was most likely later on incorrectly assigned to that taxon, or the genome KF024722 incorrectly assigned to that species. KF024722 is indeed nearly identical to the *Mycobacterium phage rosebush* (genus *Pgonelikevirus*) genome AY129334 and maximally supported as its sister group (Supplementary File S3).

Similarly, the species *Enterobacteria phage T5* appears as non-monophyletic in the tree because of the positioning of genome AY509815 (Supplementary Files S1, S3). Its genome size clearly deviates from the main group of three T5 genomes, caused by AY509815 being incomplete but aberrantly annotated. The GBDP formulas selected as optimal (Table 1) presuppose complete genomes, whereas, e.g., formula *d*_4_ (Meier-Kolthoff et al. 2013) placed all four T5 genomes in a well-supported monophyletic group (data not shown).

### Clusters vs. classification

The tree includes groups of very closely related phage species that would form a single species according to the optimal species delineation threshold, such as the nearly identical *Enterobacteria phage f1*, *fd* and *M13*; *Caulobacter phage karma*, *magneto*, *phicbk* and *swift*; *Mycobacterium phage arturo*, *backyardigan*, *LHTSCC* and *peaches*, *Pseudomonas phage 14-1*, *F8*, *LBL*, *LMA2*, *PB1*, and *SN*; and *Staphylococcus phage K* and *G*(Supplementary File S3). Whereas these clusters corresponded to reasonably to maximally (83%, 100%, 100%, 100%, 100%) supported clades, branch support for the assigned species was lacking, probably because of the high uniformity of the genomes within the clusters. Conversely, a number of species were suggested to be split. For instance, *Vibrio phage CTX* was distributed over four clusters, in agreement with differences in genome size, whereas *Enterobacteria phage P1* and *Enterobacteria phage Qbeta* were split into two clusters, respectively. Dissecting such species did not necessarily yield higher branch support though.

Overall, however, taxon support for the taxon-derived clusters was considerably higher than for the taxa themselves at the genus and species rank when inferred from amino-acid sequences (Table 1, Figure 3). A similar but weaker effect was observed for the nucleotide sequences, whereas no improvement was observed at the level of families. This not only indicated that deviations from ultrametricity hardly present a problem for the selected methods but also that, except for the families, from a phylogenetic viewpoint the classification would benefit from modifications that yielded taxa of more uniform genetic divergence. The MRI value for the genera was considerably larger than for the species, hence more effort would be needed to transform the species into groups of comparable divergence (Table 1).

**Figure 3.**
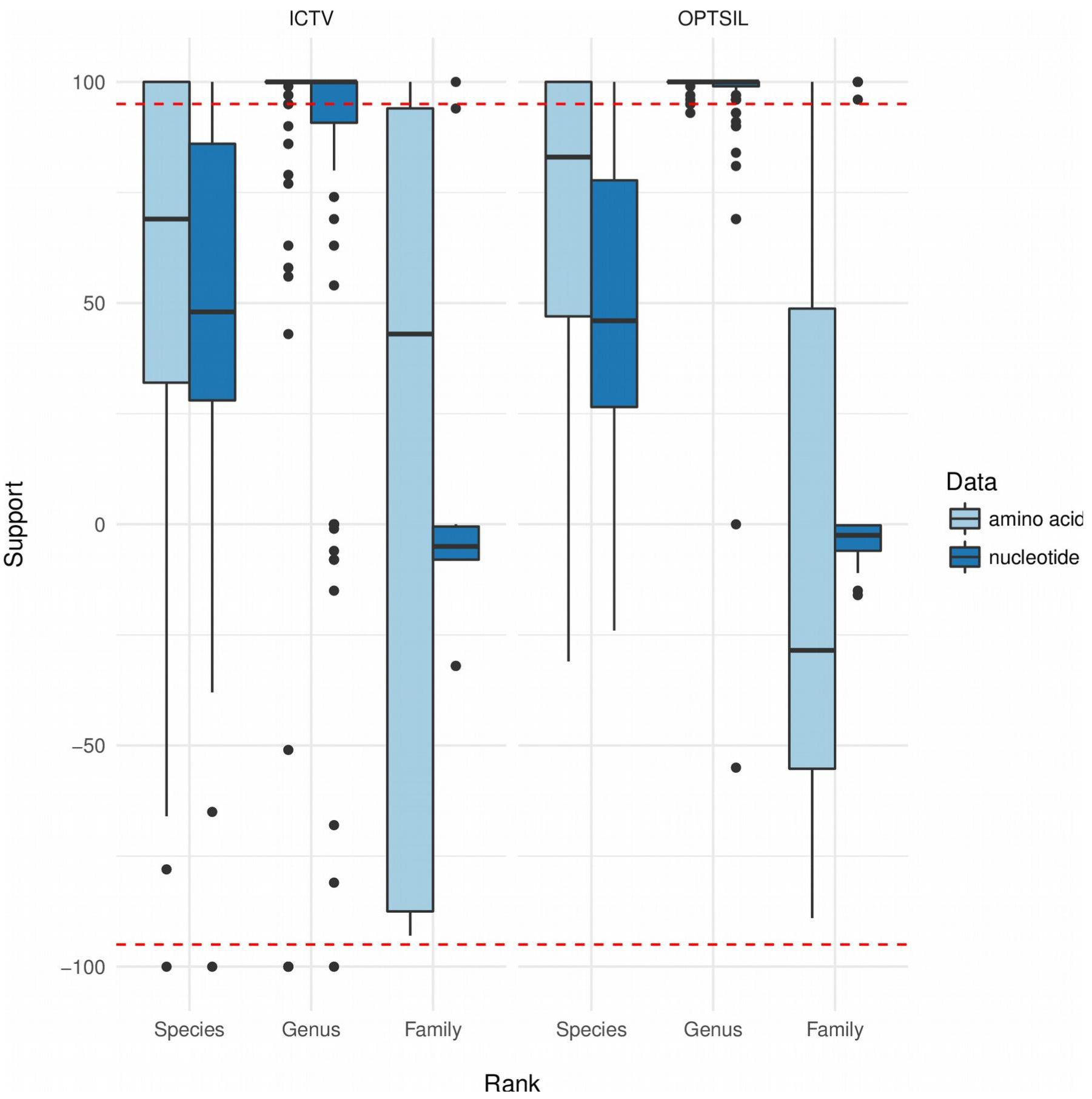
Taxon support at each taxonomic rank for ICTV taxa and clusters derived from these taxa. The underlying phylogenies and clusterings are based onthe optimal GBDP settings provided in Table 1 for nucleotide and amino-acid sequences, the clusterings also on the according distance thresholds T. The dashed lines indicate areas of significant support ( ≥ 95%) or conflict ( ≤ −95%); taxon support is displayed as negative in the case of non-monophyletic taxa.

**Table 1:** Optimal GBDP settings. The best GBDP settings according to the two-step Pareto multi-objective optimization based on (1) taxon support and (2) Modified Rand Index (MRI) at each taxonomic rank, after optimizing the distance threshold T separately for each rank and an F value of 0.5. Numeric results for the ICTV reference data sets are also provided.

|  | amino acid | nucleotide |
| --- | --- | --- |
| Word length | 3 | 11 |
| E-value filter | 0.1 | 1.0 |
| Algorithm | greedy-with-trimming | greedy-with-trimming |
| Formula | D6 | D0 |
| Average support | 56.52 % | 36.77 % |
| Number of species | 480 | 480 (same as left col.) |
| Number of genera | 98 | 98 (same as left col.) |
| Number of families | 15 | 15 (same as left col.) |
| Taxon support, species | 0.87 | 0.90 |
| Taxon support, genus | 0.95 | 0.96 |
| Taxon support, family | 0.47 | 0.73 |
| MRI, species | 0.67 | 0.67 |
| MRI, genus | 0.89 | 0.89 |
| MRI, family | 0.72 | 0.34 |
| T, species | 0.118980 | 0.022085 |
| T, genus | 0.749680 | 0.842700 |
| T, family | 0.985225 | 0.997455 |
| Number of species clusters | 387 | 474 |
| Number of genus clusters | 115 | 132 |
| Number of family clusters | 10 | 32 |
| Taxon support, species clusters | 0.99 | 0.98 |
| Taxon support, genus clusters | 1.00 | 0.99 |
| Taxon support, family clusters | 0.42 | 0.82 |
| Species to be split | 2.30 % | 2.50 % |
| Genera to be split | 9.10 % | 16.20 % |
| Families to be split | 33.30 % | 40.00 % |
| Multi-species clusters | 17.10 % | 4.60 % |
| Multi-genus clusters | 7.80 % | 4.50 % |
| Multi-family clusters | 70.00 % | 37.50 % |

### Analysis of the expanded phage data set

The comprehensive data set contained 4419 genomes, which amounted to 9,765,990 pairwise comparisons. Supplementary File S5 shows the amino acid-based phylogenomic tree inferred from the amino-acid sequences of the 4K genomes under the best GBDP setting. The application of the optimal *T* values and the OPTSIL program yielded an assignment into 49 families, 721 genera and 2511 species. These differences from the reference data set (Table 1) are as expected given the c. seven times larger expanded data set. The distribution of taxon support in the expanded data set (Supplementary File S5) was similar to the one for the reference data set (Fig. 3). A total of 2116 new species clusters (84%), i.e. clusters not covering any species listed in the ICTV classification, 603 new genus clusters (83%) and 35 new family clusters (57%) were found. The three largest new species clusters comprised mainly *Enterobacteria phage phiX174* (98 genomes), mainly *Propionibacterium phage*(88 genomes) and mainly *Synechococcus phage ACG-2014d* (45 genomes), respectively.

The plots of the host specificity in Fig. 4A do not indicate a marked difference between the ICTV classification, the clustering of the reference data set and the clustering of the expanded data set. Most phage species appeared to be specific at the level of host species, and there was no overall trend of decreasing host specificity with increased sampling. Phage species not specific for a single host species were mostly specific for a host genus and, with a single exception, specific for a host family (Fig. 4B). Entire phage genera were not in general specific for host species, and even specificity for host genera decreased for better sampled taxa; as a rule, phage genera were specific for host families. Phage families did not display any specificity that was independent of sampling (Fig. 4B).

**Figure 4.**
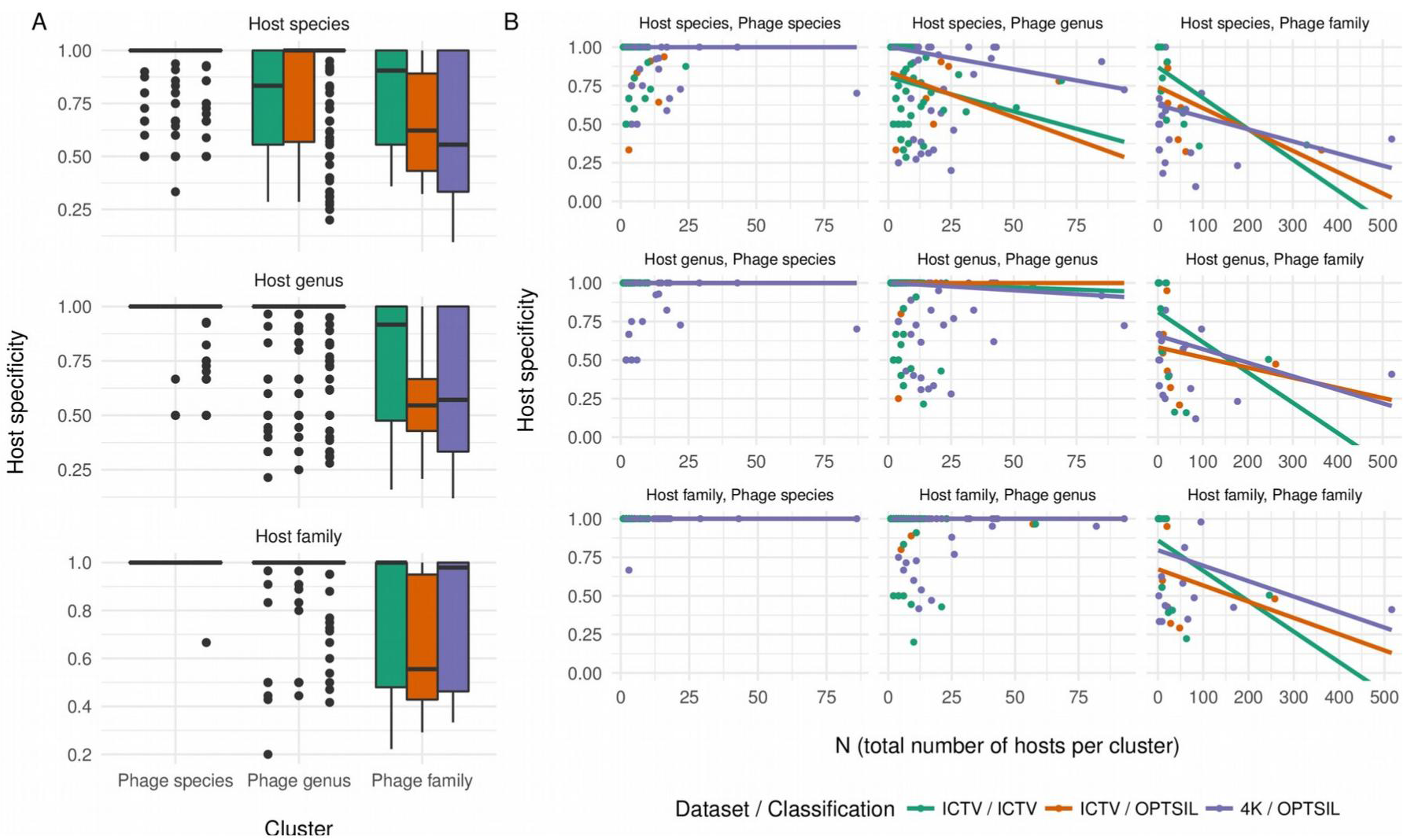
Host specificity at all taxonomic ranks. Panel A shows the distribution ofthe specificity at each of the three taxonomic ranks investigated for host taxa on the one hand and ICTV taxa as well as clusters derived from these taxa (Table 1) on the other hand. Panel B shows the host specificity additionally in dependence on N, the overall number of phages with an interpretable host entry per taxon or cluster, as visualized using separate robust-line fits for each subset of the data. Clusters were calculated from both the reference (ICTV) and the expanded (4K) data set.

## Discussion

### GBDP for classification and phylogeny of phages

Our results clearly indicate that phage phylogeny and classification with GBDP is feasible. Except for the family rank, the selected settings yielded reasonably supported trees that agreed well with G+C content values and genome sizes (Fig. 2). Only a handful of significantly supported conflicts with the ICTV classification remained, which were uniformly due to misclassifications or incorrectly annotated GenBank sequences. This success might be due to the great adaptability of GBDP to solving specific phylogenomic questions, as well-performing settings can be chosen from a large number of distinct parameter combinations, which appear to cover a range of options comparable to if not exceeding the one known from the phage literature (Supplementary File S1).

Moreover, the optimized clustering approaches, even though they are not proper phylogenetic methods and are potentially affected by deviations from a molecular clock, here yielded higher taxon support than the original ICTV taxa, particularly at the genus rank and at the species level in the case of amino-acid sequences. This holds even though the inferred phage phylogeny partially strongly deviates from a molecular clock (Fig. 2). Whereas earlier studies (Krupovic et al. 2016) reported “spurious taxonomic lumping” when applying (potentially non-optimal) clustering methods to phage genomes, we observed on average higher branch support for the clusters rather than the original taxa. Thus parts of the ICTV classification could be improved phylogenetically, too, by generating some phage species and phage genera more uniform in terms of sequence divergence. Whereas we believe that this would ease the interpretation of phage classification as in the case of bacterial species (Auch et al. 2010a; Meier-Kolthoff et al. 2013) and would not affect the host specificity of the resulting taxa (Fig. 4), our primary criterion was phylogenetic support for the current taxa. It is up to the phage taxonomist to which extent sequence divergence should be used as secondary criterion, but augmenting GBDP phylogenies with clustering results is likely to assist in delineating phage taxa. The ideal classification would then only contain phylogenetically well-supported taxa displaying a sequence divergence within the range typical for their rank. Apparently such classification could realistically be obtained under optimal GBDP settings at the species and genus rank.

Distinct structural annotations of a phage genome might yield distinct numbers of protein sequences. A nucleotide sequence of a phage might even represent overlapping genes and code for multiple proteins (Chirico et al. 2010). The optimal GBDP settings include formula *d*_0_ or *d*_6_ (Table 1), which consider gene content (Meier-Kolthoff et al. 2013), and could thus be vulnerable against differences in protein composition due to distinct annotation. However, part of these issues might already be removed by the culling of paralogous sequences conducted by GBDP (Henz et al. 2005), and we did not observe any apparently annotation-related issues in our analyses (Fig. 2). Moreover, GBDP works well with nucleotide sequences at the level of phage species and genera (Fig. 3), which helps avoiding annotation artifacts entirely.

Phages are known to be affected by horizontal gene transfer (HGT), and, particularly when occupying similar ecological niches, this can lead to a high degree of mosaic diversity (Hendrix 2003). However, the extent of HGT varies between different virus families and only partially affects certain viral functions (Krupovic et al. 2011). Whereas some authors have concluded that hierarchical classification should rather not be aimed at in such situations (Nelson 2004; Hatfull 2008; Lima-Mendez et al. 2008), we believe that as in the case of Archaea and Bacteria, the Linnaean hierarchy is feasible and useful despite a prevalence of HGT (Klenk and Göker 2010). Rather, phylogenetic inference should ensure that branch support reflects the proportion of genes in agreement with that branch. A taxonomic classification that assigned taxa only to well-supported clades could then hardly be called into question because of HGT. The partition bootstrap, which bootstraps entire genes instead of single positions in concatenated gene alignments has been suggested to reduce conflict and to provide more realistic support values in phylogenomic analyses (Siddall 2010). The here selected best GBDP methods use pseudo-bootstrapping in conjunction with the greedy-with-trimming algorithm (Meier-Kolthoff et al. 2014a), which is as close as possible within the GBDP framework to the partition bootstrap (Hahnke et al. 2016). Hence, a strong GBDP pseudo-bootstrap value for a branch indicated that it is supported by at least the majority of the genes.

Although a common evolutionary origin of *Caudovirales* has been proposed (Veesler and Cambillau 2011) and further studies suggested that most of the genes of contemporary phages derive from a common ancestral pool of genes (Hendrix et al. 1999), multiple origins of phages might argue against a phylogenetic framework for their classification and even against their comprehensive phylogenetic analysis (Fig. 2). However, a carefully chosen method for the inference of trees, when applied to several lineages of independent origin, should simply leave their relative positioning, i.e. the backbone of the tree, unresolved. A taxonomic classification focusing on well-supported branches could then hardly be affected by the independent origin of the major lineages.

Lack of support at the backbone of the phylogenetic tree (Fig. 2) and for the phage families (Fig. 3) was indeed been observed for the current data set, and could not be solved by switching to clustering approaches optimized for the family rank. Rampant early HGT and multiple origins of phages are possible causes of this lack of resolution. Still stronger support for the phage families at the amino-acid level rather than the nucleotide level (Fig. 3) alternatively points to a fading of the phylogenetic signal due to the rapid evolution or an ancient origin of phages (Hatfull 2008), in line with the fact that affiliation of phages to the same family is currently possible without any DNA sequence relatedness (Nelson 2004). Within the current GBDP framework, this issue could only be solved by creating less divergent phage families. We do not think such a step is necessary, however, since the assignment of phages to families is currently mainly based on characteristics such as morphology, replication mode and overall genomic architecture (Hendrix et al. 1999); some of these features, such as neck organization of tailed phages, can even be inferred from sequences (Lopes et al. 2014). Conversely, convergent evolution can explain the resemblance between apparently unrelated viruses (Ackermann 2007), and we cannot rule out that some of the character states used to define phage families are plesiomorphic or homoplasious (Hennig 1965; Farris 1974; Wiley and Lieberman 2011) and in conflict with sequence data (Nelson 2004). However, GBDP is not suggested as a replacement for these features but for solving questions their analysis cannot address. The currently hardly applied subfamily rank could be introduced in the future for phylogenomically well supported groups of genera below the level of the family.

The analysis of the expanded data set demonstrated that the optimized GBDP settings can be used to classify phages at the level of species and genera, even phages not yet listed in the ICTV taxonomy. We did not observe significant differences to the reference data set in terms of the quantitative behavior of the taxa at the distinct ranks. Rather, the vast majority of potential taxa are simply not yet covered by the ICTV classification (Roux et al. 2015). We thus expect an increased taxonomic coverage to yield little conflict but new insights due to the sheer amount of data.

For instance, the comparison of the reference and the expanded data set indicated that the specificity of phage species for host species and that of phage genera for host families is not an artifact of insufficient sampling but rather a real feature of the data. Switching from the ICTV taxa to OPTSIL clusters derived from these taxa and increasing the genome sampling to the expanded data set did not change these patterns. Phages observed in the laboratory are indeed mostly specific to host species, and the analysis of marine phages even indicates strain specificity in some cases (Chibani-Chennoufi
et al. 2004). In contrast, broad host ranges are rather uncommon for phages; such polyvalence which might be an adaptation to low concentrations of host cells (Chibani-Chennoufi
et al. 2004). Even though within the scope of this study we could not account for possibly still biased sampling, for wrongly assigned phage hosts and for prokaryotic taxa that do not reflect the phylogenetic relationships of the host, the results on host specificity thus appear entirely reasonable.

## Publicly available web service

The best combination of GBDP parameters determined in this study for phylogenetic inference from whole nucleotide and proteome sequences of phages were incorporated into the standalone web service “VICTOR”, the VIrus Classification and Tree building Online Resource. Its results include phylogenomic trees with branch support and augmented with suggestions for taxon boundaries, thus allowing for an informed taxonomic decision. Users can specify sequences of up to 50 phages in several file formats as well as indicate whether nucleotide or amino-acid sequences should be analyzed. When complete genome sequences are missing for some phages of interests, users can refrain from considering the optimal formulas *d*_0_ or *d*_6_ (Table 1) and use the formula *d*_4_ instead, which is not the best performing formula when applied to completely sequenced genomes but robust against the use of incomplete ones (Meier-Kolthoff et al. 2013).

The web service is available as a rapid yet reliable bioinformatics application free of charge at http://ggdc.dsmz.de/victor.php. Whereas it was beyond the scope of our study to address groups of viruses other than phages, we see no reason why the service should not be used to elucidate the phylogeny of plant and animal viruses, and to determine how to best delineate their taxa with GBDP is a logical next step.

## Materials and Methods

### Reference data set

Genomes were taxonomically selected by querying the INSDC databases for all species names assigned to phage families in the third version of the 2015 ICTV master species list (King et al. 2012), which contained a total of 548 species, 104 genera and 18 families; we did not observe a new version in 2016 that contained more taxa. Using all available whole-genome sequences of phages instead would enrich the data set with informal taxon names that could hardly be compared to each other and to the formally accepted names in the ICTV master list. Genomes assigned to species *sensu lato* were also removed. The collected data were further restricted to complete genome sequences containing protein annotation. Duplicate genomes (due to distinct annotation versions) were detected using MD5 checksums (Rivest 1992) calculated from their nucleotide sequences and only the version with most protein sequences kept. The reference data set is listed in Supplementary File S1.

### Distance calculation

Pairwise intergenomic distances were calculated between amino acid and nucleotide sequences with the current version of the Genome BLAST Distance Phylogeny (GBDP) approach (Meier-Kolthoff et al. 2013), including 100 pseudo-bootstrap replicates (Meier-Kolthoff et al. 2014a) for calculating branch support. BLAST+ (Camacho et al. 2009) was used as local alignment tool under default settings but with a broad range of e-value filters (10, 1, 1e-1, 1e-2, 1e-3, 1e-8). GBDP was run with two distinct algorithms (trimming and coverage) for filtering the BLAST+ output as well as its ten distance formulas (Meier-Kolthoff et al. 2013)for exploring either gene content or sequence identity or both. Thus a total of 240 unique combinations of parameters were investigated (two types of sequence dat six e-value settings × two GBDP algorithms × ten GBDP distance formulas). All settings are included in Supplementary File S1 together with the respective results.

### Inference and assessment of phylogenetic trees

Phylogenetic trees were inferred from the original and pseudo-bootstrapped distance matrices using FastME 2.1.4 (Lefort et al. 2015) and SPR branch swapping (Farris 1972) and rooted using the midpoint method (Hess and De Moraes Russo 2007). FastME is topologically more accurate than the well-known neighbour-joining (NJ) algorithm (Saitou and Nei 1987) because both are based on the balanced minimum evolution criterion, which NJ only greedily optimizes (Lefort et al. 2015). Checks for monophyletic, paraphyletic and polyphyletic taxa were conducted based on the criteria of Farris (Farris 1974) and in-house developed scripts (Hahnke et al. 2016). The bootstrap support of each non-trivially monophyletic taxon was recorded as well as the support against each non-monophyletic taxon as the maximum support among all clades in the rooted phylogeny that were in conflict with the monophyly of that taxon. The sum of the support values for the taxa at each taxonomic rank, relative to the sum of all support values, either for or against the monophyly of a taxon, yielded a measure of overall phylogenetic fit (“taxon support”) of a given tree, and thus of the underlying distance matrix and combination of GBDP parameters, to the taxonomic classification at that rank. If most taxa of a certain taxonomic rank in the ICTV classification (King et al. 2012) received little positive or negative support, this would indicate insufficient resolution of the respective method at that rank, whereas strong support against many taxa would indicated a bias of the method. In contrast, single taxa with strong support against their monophyly rather pointed to a problem with the classification or just with the naming of certain INSDC entries.

### Inference and assessment of clusterings

In clustering experiments additional to phylogenetic inference, the OPTSIL software (Göker et al. 2009) was used to optimize distance thresholds *T* by maximizing the agreement of the resulting non-hierarchical clustering with a given reference partition. Agreement was quantified as the modified Rand index (MRI) (Göker et al. 2009), which is equal to 1 in the case of two identical partitions (Hubert and Arabie 1985) and proportionally lower depending on the amount of disagreement. The ICTV classification reduced to each rank was used as reference partition; genomes from species not assigned to a genus were removed when analyzing the genus rank. The *F* parameter of OPTSIL was set to 0.5, which yielded the highest clustering consistency in earlier studies (Meier-Kolthoff et al. 2014b). The fit of each distance matrix, and thus each combination of GBDP parameters, to the taxonomic classification at each rank can then be represented as the highest obtained MRI value. However, clusters inferred from the same distance matrix as a phylogenetic tree that yielded strong support against their monophyly would point to problems due to non-ultrametricity in the data (Meier-Kolthoff et al. 2014c). For this reason, taxon support for the optimal clusters was determined as described above.

### Correspondence between criteria and optimal settings

Since each taxonomic rank yielded a separate optimality criterion regarding both taxon support and MRI, the correspondence between the six optimality criteria was explored with a principal-components analysis as implemented in the FactoMineR package (Lê et al. 2008) for the R statistical environment (R Development Core Team 2015). A Pareto multi-objective selection was conducted with the rPref package (Roocks 2016) for R (R Development Core Team 2015) to determine the subset of equally feasible choices ("Pareto frontier") for nucleotide and amino-acid sequences, respectively. The taxon support for each of the three ranks family, genus and species served as the first set of objectives; the resulting subset of GBDP settings was further reduced using the MRI values as second set of objectives. R code to analyze and plot the optimality criteria is provided in Supplementary File S2.

### Analysis of expanded data set

A larger data set of 4419 phages was collected from GenBank and the PhAnToMe FTP server (as of July 2016) (Overbeek et al. 2014), without restriction to taxa recognized by the ICTV (Supplementary File S1). Using the best GBDP settings for the analysis of amino-acid sequences and the thresholds for delineation at the species, genus and family rank, phage diversity was quantified and compared to the one of the reference data set. In addition to the number of new and already known clusters or taxa and the number of genomes per cluster or taxon, the effect of increased genome sampling on host specificity was examined. The “specific host” entry was extracted from the GenBank files and restricted to validly published names of host taxa as listed in Prokaryotic Nomenclature Up-To-Date (October 2016, https://www.dsmz.de/bacterial-diversity/prokaryotic-nomenclature-up-to-date.html). The thus standardized host names were used as-is; fixing prokaryotic taxa that do not reflect their phylogenetic relationships (Klenk and Göker 2010) was beyond the scope of the present study. Finally, the host specificity of each cluster was assessed in analogy to the Berger-Parker-Index (Berger and Parker 1970) via the formula *m*/*N* with *m* being the frequency of the most frequent host and *N* the total number of (potentially) hosts indicated for the genomes in that cluster. The dependency of this index on *N* was studied using robust line fits as implemented in R (R Development Core Team 2015) since high specificity might be a sampling artifact.

## Contributions

JPMK and MG devised the study. JPMK performed the experiments, analyzed the data, conducted the (statistical) analyses and implemented the web service. JPMK and MG wrote the manuscript.

## Acknowledgements

We thank Johannes Wittmann (DSMZ) for drawing our attention to the need for genome sequence-based methods in virus taxonomy. We thank Marek Dynowski (formerly Zentrum für Datenverarbeitung, University of Tübingen; now: University of Manchester) and Werner Dilling (Zentrum für Datenverarbeitung, University of Tübingen) for granting access to the bwGRiD computing infrastructure and for their technical support, which enabled us to conduct parts of the high-throughput computations on the bwGRiD. This work was supported by Deutsche Forschungsgemeinschaft within TRR 51.

### Supplementary Material

- S1 Supplementary File S1 – First sheet, list of INSDC accessions of (annotated) genome sequence that were successfully mapped to a phage species from the ICTV master table, G+C content values, genome sizes, complete ICTV annotation and cluster affiliation under the selected GBDP settings. Second sheet, taxon support and MRI values obtained for all 240 examined GBDP settings. Third sheet, phylogenetic results for ICTV taxa obtained using amino-acid sequences, assessed under the respective optimal GBDP settings. Fourth sheet, phylogenetic results for ICTV taxa obtained using nucleotide sequences, also assessed under the respective optimal GBDP settings. Fifth sheet, phylogenetic results for OPTSIL clusters inferred from the amino-acid sequences. Sixth sheet, phylogenetic results for OPTSIL clusters inferred from the nucleotide sequences. Seventh sheet, list of phage genomes within the expanded data set, G+C content values, genome sizes and cluster affiliation under the selected GBDP settings. Eighth sheet, cross-comparison of tools for the sequence-based comparison of phages, illustrating the specifics and limitations of each methods.
- S2 Supplementary File S2 - R code used for the statistical analyses, given file S1.
- S3 Supplementary Table S3 - Phylogenomic tree in linear representation inferred from the amino-acid sequences of the reference data set under optimal GBDP settings.
- S4 Supplementary File S4 - Phylogenomic tree inferred from the nucleotide sequences of the reference data set under optimal GBDP settings.
- S5 Supplementary File S5 - Phylogenomic tree inferred from the amino-acid sequences of the expanded data set under optimal GBDP settings. Also contains a plot of the taxon support of the inferred clusters at each rank.

